# Timed Notch Inhibition drives Photoreceptor fate specification in Human Retinal Organoids

**DOI:** 10.1101/2022.05.19.492679

**Authors:** Shereen H. Chew, Cassandra Martinez, Kathleen R. Chirco, Sangeetha Kandoi, Deepak A. Lamba

## Abstract

**Purpose:** Transplanting photoreceptors from human pluripotent stem cell derived retinal organoids have the potential to reverse vision loss in affected individuals. However, transplantable photoreceptors are only a subset of all cells in the organoids. Hence the goal of our current study was to accelerate and synchronize photoreceptor differentiation in retinal organoids by inhibiting the Notch signaling pathway at different developmental time-points using a small molecule, PF-03084014 (PF).

**Methods:** Human induced pluripotent stem cell (hiPSC)- and embryonic stem cells (hESC)-derived retinal organoids were treated with 10μM PF for three days at day 45 (D45), D60, D90 and D120 of differentiation. Organoids collected at 14-, 28-, and 42-days post-PF treatment were analyzed for progenitor and photoreceptor markers and Notch pathway inhibition by immunohistochemistry (IHC), quantitative PCR (qPCR) and bulk RNA-seq (n=3-5 organoids from 3 independent experiments).

**Results:** Retinal organoids collected at 14-days post-PF treatment showed a decrease in progenitor markers (KI67, VSX2, PAX6, and LHX2) and an increase in differentiated pan-photoreceptor markers (OTX2, CRX, and RCVRN) at all organoid stages except D120. PF-treated organoids at D45 and D60 exhibited an increase in cone photoreceptor markers (RXRG and ARR3). PF-treatment at D90 revealed an increase in cone and rod photoreceptors markers (ARR3, NRL, and NR2E3). Bulk RNA-seq analysis mirrored the IHC data and qPCR confirmed Notch effector inhibition.

**Conclusions:** Timing the Notch pathway inhibition in human retinal organoids to align with progenitor competency stages can yield an enriched population of early cone or rod photoreceptors.

## INTRODUCTION

Most forms of retinal degeneration ultimately lead to the permanent loss of the light-sensing cells called photoreceptors. The human retina does not inherently regenerate and so the ensuing vision loss is permanent. Currently, there are no known effective therapeutics for vast majority of these patients. However, as long as the inner retina stays intact, transplanting photoreceptors can provide an avenue to restore vision. Human pluripotent stem cells (hPSCs) have been utilized to generate retinal photoreceptors in a monolayer or organoid setting to study development and disease.^1–4^ Alternatively, lab-generated photoreceptors could be used as a pool of photoreceptors for transplantation.^5–8^ Over the course of development, retinal progenitor cells (RPCs) differentiate into their final cell fate in the neural retina, including cone and rod photoreceptors (PRs), retinal ganglion cells (RGCs), amacrine cells (ACs), horizontal cells (HCs), bipolar cells (BCs), and Müller glia (MG).

One of the critical pathways maintaining RPC multipotency is the Notch pathway. It is a ligand-receptor pathway that is used between multipotent cells to control neuronal potential and proliferation.^9,10^ When the ligand (Delta-like or Jagged in mammals) from a neighboring cell interacts with the Notch receptor, it induces cleavage in Notch by ADAM-family metalloproteases (Figure 1A).^10^ The Notch intracellular domain (NICD) is then cleaved by γ-secretase and activates the DNA-binding CSL protein, which complexes with Mastermind and other transcription factors, in the nucleus to express proliferation genes, such as *HES1* and *HES5* (Figure 1A).^10^ This process helps to maintain the RPC pool throughout retinal development. Notch signaling have been studied in many species to understand its importance in retinal differentiation and specification. Constitutive Notch pathway activation results in either a persistent progenitor state or differentiation into MG.^11^ Conversely, inhibition of Notch signaling via small molecule γ-secretase inhibitors in mouse and chicken retina leads to synchronized differentiation of the tissue.^12^ The Notch signaling pathway has also been implicated in retinal regeneration: in *Xenopus* and chicken retina, Notch up-regulates MG following damage and is critical to restore the progenitor potential.^13,14^

**Figure 1.**
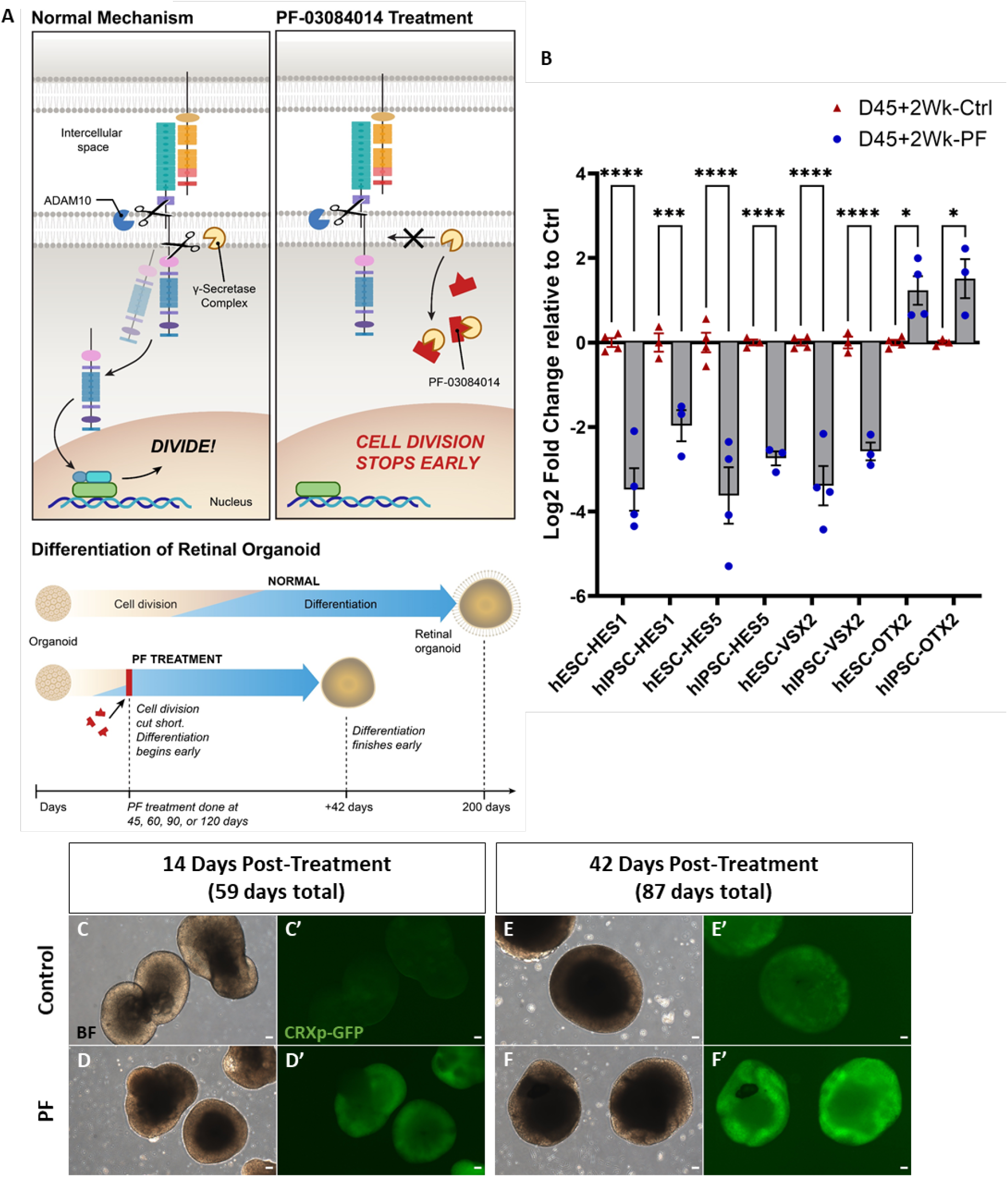
Retina progenitor cells lose Notch activity after PF treatment and increase in photoreceptor fate. (A) Schematic of the Notch Pathway before and after treatment with PF. (B) qPCR data are shown for hESC and hiPSC lines after 14 days post-treatment as log2 fold change compared to control organoids for *HES1, HES5, VSX2*, and *OTX2* (n = 5 organoids per line). (C-F’) Bright-field and GFP expression observation in early retinal organoid generated from CRXp-GFP H9 hESCs. PF treatment started at D45. After 14 days, PF-treated (D’) organoids expressed more CRXp-GFP than their control counterparts (C’). This was also the case after 42 days (E’, F’). Scale bar, 50μm. *p < 0.05, **p < 0.01, ***p < 0.005, ****p < 0.001. All statistical analyses were performed using one-way ANOVA with a Dunnett test to correct for multiple comparisons in GraphPad Prism 8 software.

Dysregulated Notch signaling has been associated with a number of developmental and oncogenic disorders including some forms of leukemia. As a result, drug-like molecules have been developed for human therapies including γ-secretase inhibitors. PF-03084014 (PF) is a γ-secretase inhibitor used in conjunction with docetaxel in a phase I breast cancer therapy trial.^15^ Due to its potent inhibition of the Notch pathway, we used PF to drive synchronized differentiation of RPCs in our retinal organoids.^16^ Since we know approximately when each of the retinal cell types are born from birth-dating studies, we hypothesized that targeting differentiation of specific cell types by using PF would generate an enriched pool of targeted cell types. These cells could then be used for transplantation or for studying molecular pathways that guide RPC differentiation. In this report, we describe our efforts of enriching PRs from hPSC-derived retinal organoids using the Notch pathway inhibitor, PF. We show that early progenitors (before D60) are biased to generate cones while late progenitors (beyond D90) are biased towards rods.

## RESULTS

### Small molecule γ-secretase inhibitor blocks Notch pathway activity in 3D retinal organoids and drives differentiation

Using previously published 3D retinal organoid differentiation protocols,^3,4^ we generated organoids from a hiPSC line^17^ and a modified hESC line (CRXp-GFP H9^18^). Both lines consistently made retinal organoids, signified by a bright ring of retinal tissue. D45 organoids were then treated with 10μM of the γ-secretase inhibitor, PF-03084014 hydrobromide (PF), for three days and analyzed 14-days post-treatment (D59) for downstream effectors of Notch (*HES1* and *HES5*) and early retina genes *(VSX2* and *OTX2)* using qPCR. Both *HES1* and *HES5* have been shown to be downstream of Notch and are important regulators in neuronal differentiation.^19^ *VSX2* is expressed in developing RPCs, but is downregulated as cells differentiate, except in BCs and MG.^20,21^ OTX2 is expressed in PRs, BCs, and retinal pigmented epithelium.^22,23^ We observed a consistent decrease in downstream Notch pathway targets *(HES1* and *HES5)* (Figure 1B) in organoids from both cell lines following PF treatment. This was associated with a decrease in RPC genes (*VSX2*). Concomitantly, we observed an increase in the photoreceptor gene (*OTX2*) (Figure 1B). Interestingly, DAPT, another Notch γ-secretase inhibitor, which has been used to inhibit Notch in chickens and mice retinae,^12,13^ did not drive significant differentiation (data not shown). This data indicates that the clinically relevant Notch pathway inhibitor PF drives differentiation of human RPCs.

CRX is an essential transcription factor that regulates many downstream PR genes.^22–24^ The CRXp-GFP H9 reporter line, generated to report activity of *CRX* promoter in differentiating PRs, has been shown to express GFP around D37 in retinal organoids.^18^ This reporter line allowed us to estimate the timing of PR differentiation *in vitro* (Figure 1C-F). At 14- (Figure 1D’) and 42-days (Figure 1F’) post-D45 treatment, we observed significantly higher GFP expression compared to control cultures (Figure 1C’-E’) along with loss of typical bright organoid appearance on brightfield (Figure 1D, F). Thus, PF treatment drove CRX expression, suggesting that the organoids were pushed towards a PR fate along with some organoid disorganization.

### PF treated retinal organoids lose progenitor properties in early time points

Since PF treatment had a profound effect on driving differentiation in D45 retinal organoids, we next tested the effects on RPCs at different developmental stages (D45, 60, 90, and 120) and analyzed them using IHC after 14- and 42-days (Figure 2A). Overall, we observed a decrease in KI67, a pan cell cycle marker expressed in the nuclei of proliferating cells,^25^ beginning at 14-days for the D45, D60, and D90 treatment groups (Figure 2B, F, J). Interestingly, there were more KI67^+^ cells in the D60 42-days post-PF group than the D60 14-days post-PF organoids. This could be due to a few PF treatment resistant cells that continued to proliferate afterwards. Finally, both control and PF-treated D90 organoids analyzed at 42-days had little to no KI67^+^cells, while the D120 group do not have any KI67^+^ cells after 14- and 42-days (Supplementary Figure S1A), suggesting that the organoids are fully differentiated by D134.

**Figure 2.**
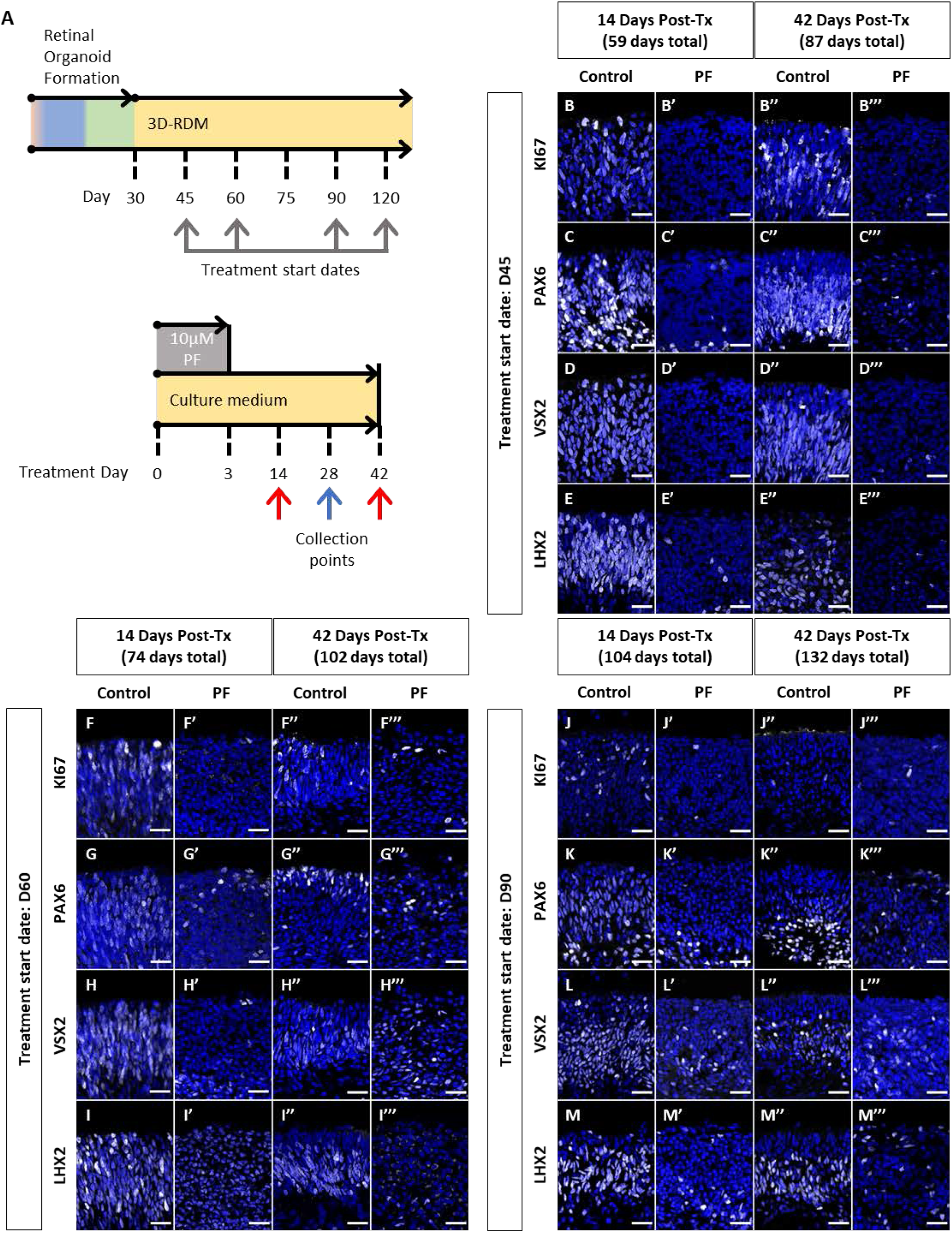
Notch inhibition turns off progenitor markers in retinal organoids. (A) Schematic of the treatment timeline. Once RPCs are generated on day 30, separate PF treatment occurred on D45, 60, 90, and 120. Collections for IHC happened after 14- and 42-days (red arrows) and for bulk RNAseq after 28-days (blue arrow). (B-M) Immunofluorescence staining using antibodies against KI67 (B, F, J), PAX6 (C, G, K), VSX2 (D, H, L), LHX2 (E, I, M) are shown in white for treatment groups D45 (B-E), D60 (F-I), and D90 (J-M). Nuclei are counterstained with DAPI in blue. Scale bar, 25μm. See also Figures S1 for D120 treatment group.

We further confirmed our KI67 data with other specific RPC progenitor markers such as PAX6, VSX2, and LHX2. We observed a consistent loss of expression of these proteins in the neuroblastic layers in the D45, D60, D90, and D120 groups after treatment (Figure 2C-E, 2G-I, 2K-M, Supplementary Figure S1B-D). Although PAX6 was expressed in the organoids, there were clear groups of PAX6^high^ and PAX6^low^ cells. These PAX6^high^ cells likely correspond with ACs, as well as RGCs but tend to die off in later stages of organoid development.^26–28^ VSX2^low^ cells were present in controls but drop to a handful after PF treatment (Figure 2C, G, K, Supplementary Figure S1B); the post treatment cells are most likely BCs, indicated by VSX2^high^ expression (Figure 2D’-D’”, H’-H’”, L’-L’”, Supplementary Figure S1C’-C’”).^21^ The same was true for remaining LHX2^+^ MG at this stage (Figure 2E, I, M, Supplementary Figure S1D).^29^ The D120 organoids contained predominantly LHX2^high^ cells (Supplementary Figure S1D), indicating that presence of few, if any RPCs, at this timepoint.

### Notch knockdown increases cone photoreceptor population in early-stage retinal progenitors

As PF treatment results in synchronized differentiation of human RPCs in the retinal organoids, we next sought to test the competency of RPCs to generate various retinal neurons. Birth-dating studies have closely correlated genesis of different retinal neurons to progenitor staging.^30–36^ Based on these, cone PRs are poised to be generated in D45-D60 retinal organoids. To investigate cone PR development, we immunostained for OTX2, CRX, RCVRN, RXRγ, and ARR3. Recoverin (RCVRN) is a Ca^2+^ binding protein that helps regulate the phosphorylation of the visual opsins in PRs.^37,38^ RXRγ is present in developing cones and is downregulated to form S-cone PRs.^18,39^ ARR3 aids in quenching the phosphorylated state of S- and M/L-opsins in cone PRs^40^. The D45 post 14-days control group has a few cells that are positive for OTX2, CRX, RCVRN, RXRγ, and ARR3 (Figure 3A-E) which increased over the next 4 weeks. Upon PF treatment, at the 14-days timepoint, we saw that the majority of cells expressed OTX2, CRX, and RXRγ along with significant increases in RCVRN (Figure 3A’-E’). At the 42-days timepoint, again we observed similar changes in the above proteins. Additionally, ARR3, which was higher at the 14-days timepoint post-treatment compared to controls, was further upregulated at 42-days post PF treatment (Figure 3E’”) suggesting further maturation of the cone PRs.

**Figure 3.**
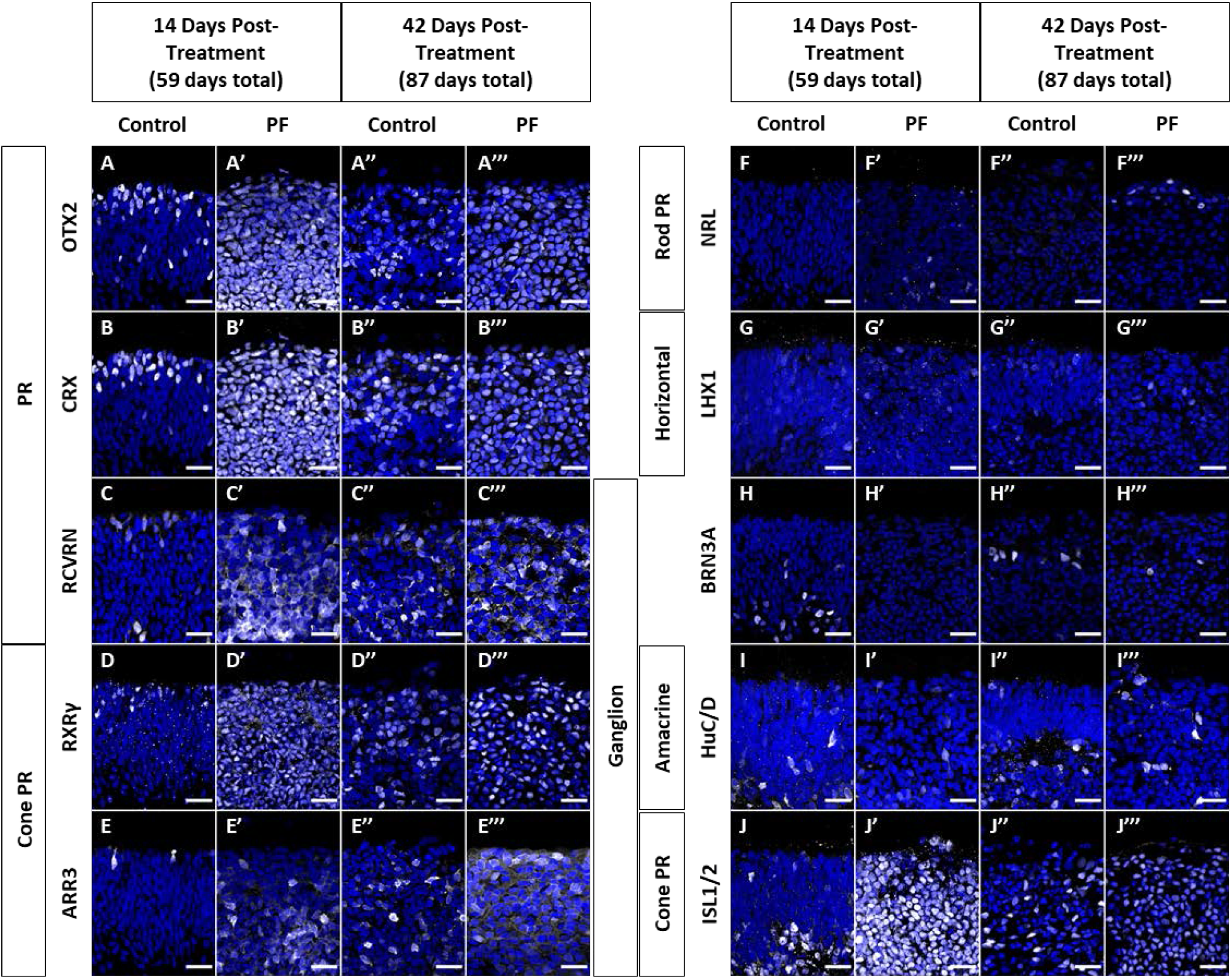
Notch inhibition at D45 causes mass generation of immature cone photoreceptors in retinal organoids after 14- and 42-days. Immunofluorescence staining using antibodies against OTX2 (A), CRX (B), RCVRN (C), RXRγ (D), ARR3 (E), NRL (F), LHX1 (G), BRN3A (H), HuC/D (I), and ISL1/2 (J) are shown in white. Markers are split into pan photoreceptors (A-C), Cone PR (D-E, J), Rod PR (F), HCs (G), RGCs (H-J), and ACs (I). Nuclei are counterstained with DAPI in blue. Scale bar, 25μm. See also Figures S2 for negative or insignificant staining.

While it is not anticipated, we wanted to assess whether D45 RPCs were competent to generate any rod PRs. NRL is necessary and sufficient to form rod PRs.^41^ We observed no NRL^+^ cells in either D45 14- and 42-days control groups (Figure 3F-F”). Surprisingly, post-PF treatment, a few rare NRL^+^ rod PRs were observed in the D45 14- and 42-days post-treated groups (Figure 3F’-F’”), suggesting even D45 retinal organoids have minor competence to generate rod PRs.

Birth-dating studies suggest that early retinal progenitors have potential to generate RGCs, ACs, and HCs.^30–33,42^ We next tested the ability to generate these cells in D45 organoids post PF-treatment. LHX1 is expressed by developing HCs,^43^ and we observed few scattered LHX1^+^HCs in control organoids (Figure 3G-G”), with no significant change upon PF treatment at both 14- and 42-days timepoints (Figure 3G’-G’”). BRN3, a POUF4 transcription factor present in RGCs in the retina, determines the diversity of RGCs.^44^ Most of the BRN3A^+^ cells are present in the lower half or below of the neuroblastic layer in the 14- and 42-days control (Figure 3H, H”).

Interestingly, upon PF treatment, we did not detect many BRN3A^+^ cells in the organoids (Figure 3H’, H’”). This finding suggests that either the RGCs are struggling to survive in a mature organoid model lacking stem or glial cell support or that PF is directly toxic to RGCs in the organoids. HuC/D is expressed by mature RGCs and ACs and in developing HCs (in rats).^45^ We found a large population of HuC/D^+^ cells in the 14-days control (Figure 3I), but the number of cells diminished modestly after treatment (Figure 3I’). The 42-days control and treated groups show similar numbers of HuC/D^+^ cells (Figure 3I”-I’”). Given the observed pattern of BRN3A^+^cells (Figure 3H), it is likely that most of the HuC/D^+^ cells are ACs in treated organoids and do not change with PF treatment. Finally, ISL1 is expressed in RGCs, ACs, and BCs while ISL2 is in cone photoreceptors.^46,47^ Staining for ISL1/2, we observed a dramatic increase in ISL1/2^+^cells in the 14-days treated group compared to the corresponding control (Figure 3J-J’) with an expression pattern similar to OTX2 and CRX staining (Figure 3A-B), suggesting that most of these might be early cone PRs. At the 42-days timepoint, there was not much change between the control and treated groups (Figure J”-J’”). This could be due to only a transient expression in ISL2 in newly differentiated cones.^48^ To test if the cellular competency changed significantly in D60 organoids, we next treated these with PF for three days and analyzed 14- and 42-days later. Here again we observed that PF resulted in a cone rich organoid (Figure 4A-E) without any significant change in the number of other cell types (Figure 4F-H).

**Figure 4.**
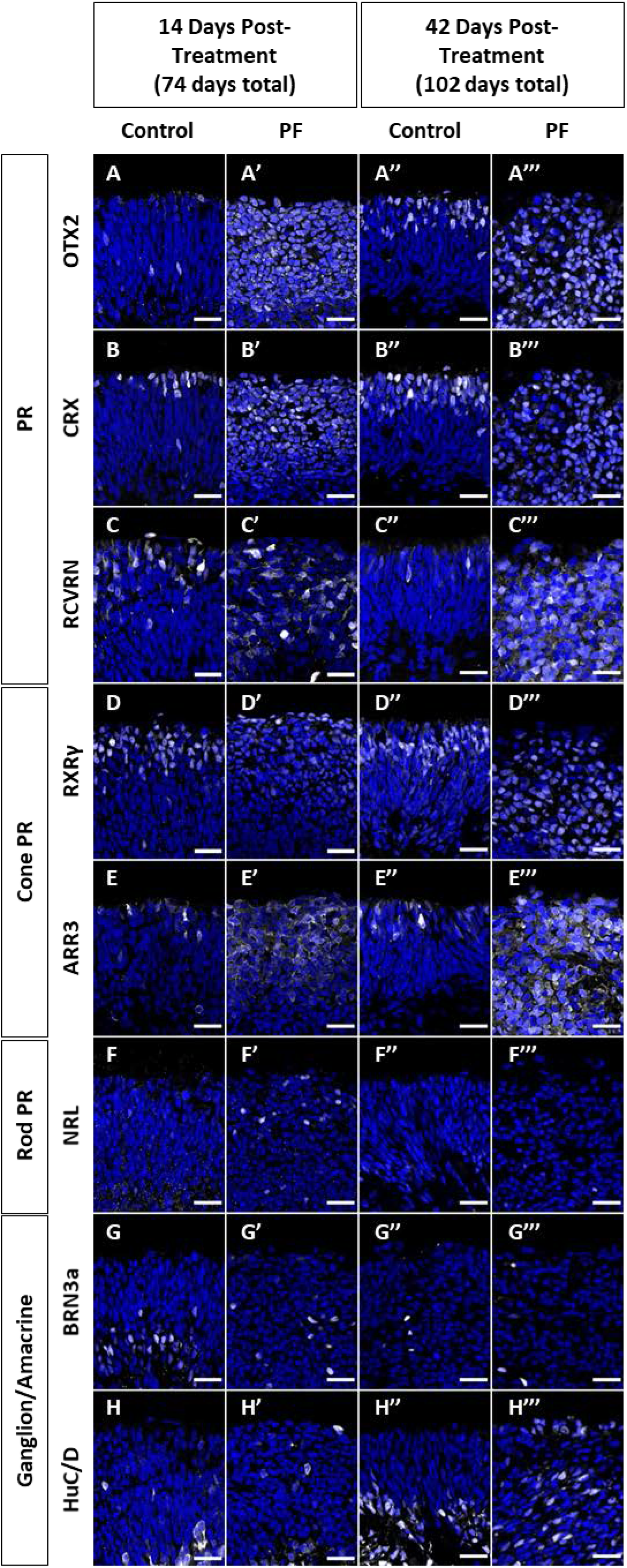
Notch inhibition at D60 causes mass generation of immature cone photoreceptors in retinal organoids after 14- and 42-days. Immunofluorescence staining using antibodies against OTX2 (A), CRX (B), RCVRN (C), RXRγ (D), ARR3 (E), NRL (F), BRN3A (G), and HuC/D (H) are shown in white. Markers are split into pan photoreceptors (A-C), Cone PR (D-E), Rod PR (F), and RGC/AC (G-I). Nuclei are counterstained with DAPI in blue. Scale bar, 25μm. See also Figures S2 for additional staining data.

One of the most intriguing findings was a lack of maturity in the cone PRs in the PF-treated differentiated organoids. S- and M/L-opsins are required for cone PRs to be able to process photons into a chemical signal. There was a lack of S- and M/L-opsins expression in both the D45 and D60 14- and 42-days post treatment groups (Supplementary Figure S2A-B, D-E). Even upon further culture for 70- and 140-days post-treatment on D45, organoids failed to express either of the opsins (Supplementary Figure 3A-B). This data suggests that while PF can drive cone-rich organoids, accelerating the *in vitro* culture system lacks the cues required to complete the maturation of PRs.

### Notch knockdown in late retinal progenitors increases rod photoreceptor population

Late retinal progenitors are biased to generate rod PRs over cones.^30–33,42,49^ To examine our ability to stimulate rod PR generation, we cultured retinal organoids for 90 days prior to a 3-day PF treatment. At 14- and 42-days post-treatment, general PR protein makers (OTX2, CRX, and RCVRN) were shown to increase in both PF-treated groups compared to control (Figure 5A-C’”). Cone PR expression also increased as shown by RXRγ and ARR3 immunolabeling (Figure 5D-E’”). Next, we looked for the generation of rod PRs using rod specific markers: NRL ^41,50^ and NR2E3 ^49,51^. We found a significant increase in both rod markers at 14-days post PF treatment and even more so after 42-days (Figure 5F-G’”). Based on these data, D90 organoids are both rod and cone PR competent. HuC/D^+^ ACs remained towards the bottom of the neuroblastic layer and did not increase following PF treatment (Figure 5H-H’”).

**Figure 5.**
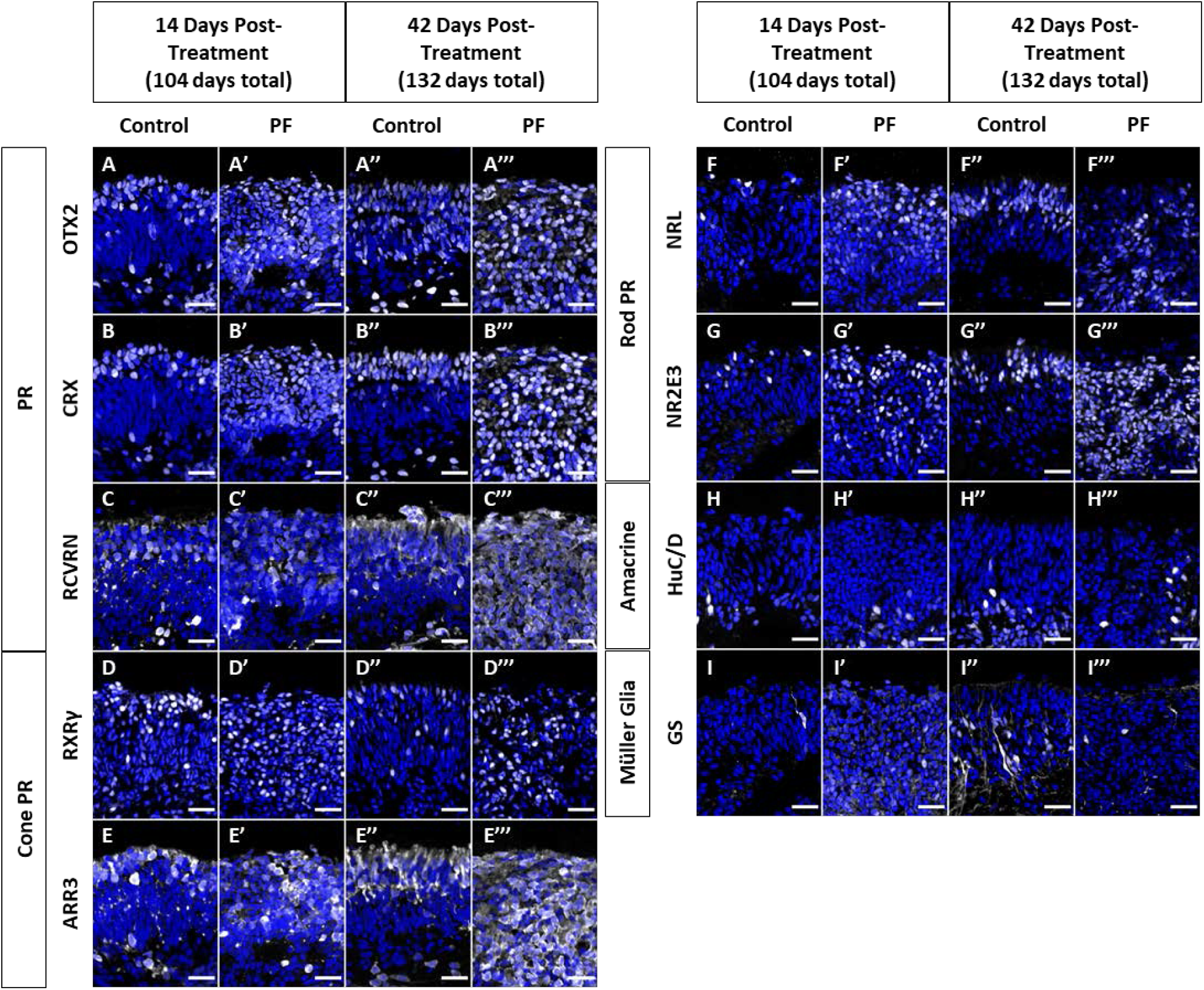
Notch inhibition at D90 causes mass generation of immature rod photoreceptors in retinal organoids after 14- and 42-days. Immunofluorescence staining using antibodies against OTX2 (A), CRX (B), RCVRN (C), RXRγ (D), ARR3 (E), NRL (F), NR2E3 (G), HuC/D (H), and GS (I) are shown in white. Markers are split into pan photoreceptors (A-C), Cone PR (D-E), Rod PR (F-G), AC/RGC (H), and MG (I). Nuclei are counterstained with DAPI in blue. Scale bar, 25μm. See also Figures S2 for additional staining data.

MG are the last retinal cell type formed from RPCs and Notch signaling is an important driver for MG formation.^52–54^ MG are identified by glutamine synthase (GS) expression.^55^ In D90 control organoids, MG start to form but are sparse (Figure 5I). Interestingly, we observed a reduction in number of GS^+^ MG in both post-treatment time points (Figure 5I’-I’”). This PF-mediated reduction in MG is most likely due to the lack of Notch signaling needed for their specification.

In the D120 treatment group, where we did not observe and change is progenitor expression following PF (Supplementary Figure S1A-D’”), there were no significant changes in expression for PR markers (OTX2, CRX, RCVRN, and ARR3; Supplementary Figure S1E-H’”). A small number of M/L-opsin^+^ cells were observed in the 42-days control and treated organoids (Supplementary Figure S1I”-I’”), but this is most likely due to the age of the organoids and not a result of the treatment. The number of NRL^+^ rod PR cells were unchanged at 14-days post-PF with a small increase at 42-days (Supplementary Figure S1J’-J’”).

### RNAseq analysis of PF-treated organoids

We next sought to analyze the effects of Notch inhibition using bulk RNAseq analysis. We cultured retinal organoids for either 45, 60 or 90 days prior to 3 days of PF treatment as above. At 28-days post-treatment, RNA was collected and processed for RNAseq analysis. This timepoint was chosen as a mid-point between the two timepoints included in the IHC analysis. Upon analysis of Notch pathway effector *(HES1* and *HES5)* expression, we observed a significant downregulation at all three stages of differentiation, with the maximal downregulation in the D45 treatment group (Figure 6A). This is likely because the organoids at this stage have most RPCs with active inter-cellular Notch activity compared to the other treatment timepoints. Similarly, we confirmed downregulation of various genes typically expressed in RPCs including *PAX6, LHX2, VSX2, ASCL1*, and *SOX2* (Figure 6A). In alignment with our IHC data, cone PR differentiation was biased in early-stage organoids (D45 and D60) as evidenced by increases in *ARR3, RXRG, TULP1, GNAT2, CNGA3*, and *PDE6C* among other cone genes (Figure 6B).

**Figure 6.**
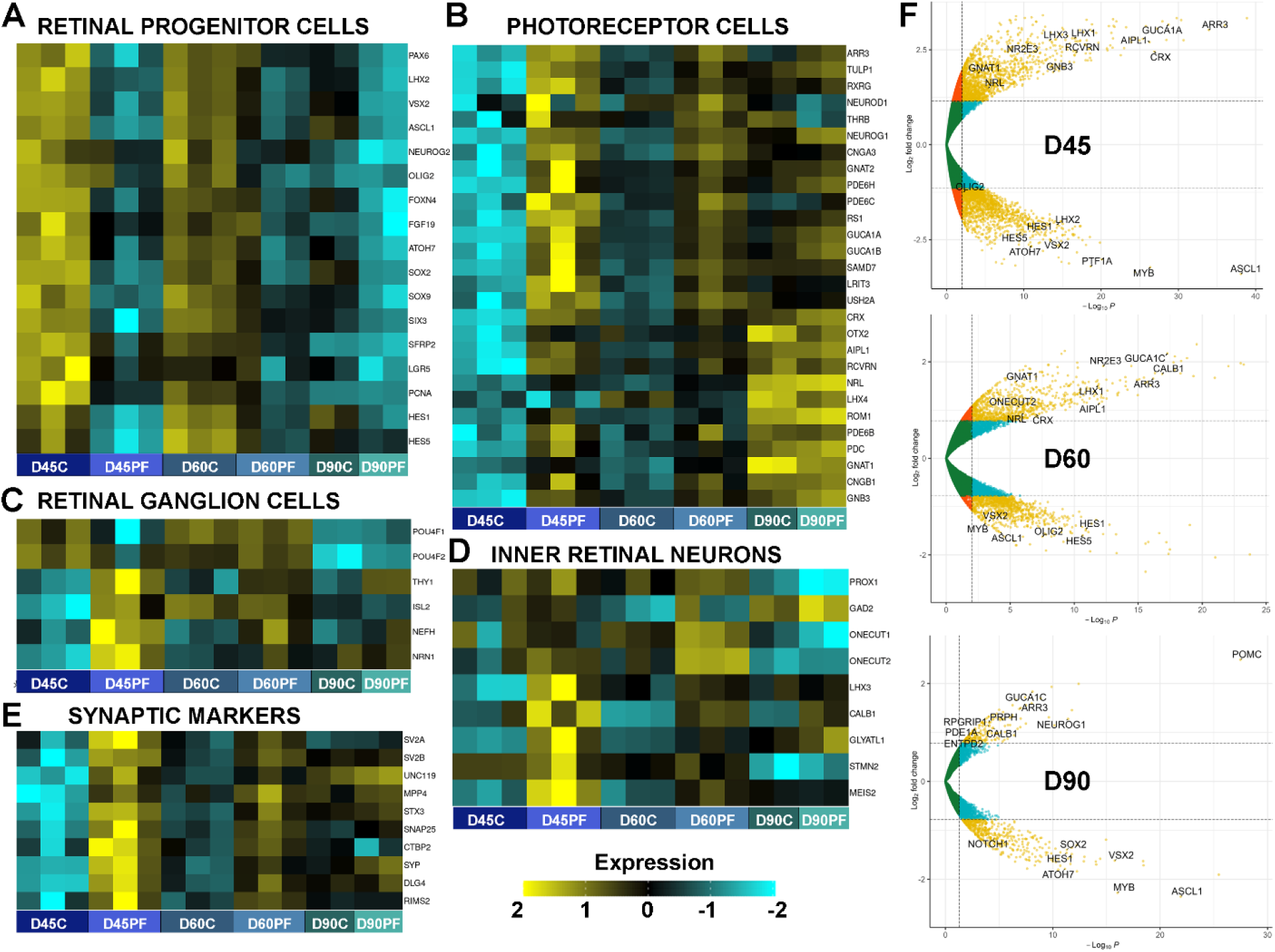
RNAseq analysis of PF treated retinal organoids confirm RPC loss and photoreceptor differentiation phenotypes. Organoids were treated for 3 days at D45, D60 and D90 of differentiation and analyzed 28-days later. (A) At all three stages there is a consistent downregulation of RPC genes and Notch pathway effectors with a stronger effect at early timepoints when the organoids are RPC enriched. (B) PR genes are highly upregulated especially at D45 and D60 treatment timepoints. Smaller increases in rod-specific genes were observed in the D90 treatment group. (C) RGC specification genes *(POU4F* family) were downregulated while maturation markers upregulated with PF. (D) Smaller increases in inner retina specification genes were observed while maturation markers were highly upregulated. (E) PF treatment led to consistent upregulation of various synaptic maturation genes expressed in the inner and outer retina. (F) Volcano plot showing the significant differentially expressed genes at all three treatment time-points.

In contrast, D90 PF-treated organoids were rod PR biased, with smaller differences in marker expression levels compared with untreated organoids at this age (D118), as RPCs have mostly differentiated at this timepoint, and control cultures have generated rods as well (Figure 6B). We observed relatively small changes in rod maturation genes *(ROM1, NRL*, and *GNB3)*.

Similar to our IHC data (Figure 3), we observed downregulation of RGC markers, *POU4F1* and *POU4F2*, in the D45 and D90 treatment groups; however, we did see an increase in maturation genes typically associated with RGCs *(THY1, NEFH*, and *NRN1)* at all three timepoints (Figure 6C). These increases in RGC markers could be due to an increase in expression in surviving RGCs, or cross-expression in other inner retinal neurons. Next, we focused on other inner retinal markers such as *PROX1*, which marks progenitors destined to interneuron fates (e.g. ACs and HCs).^56^ *PROX1* was increased in D60 PF-treated organoids, suggesting an increase in the pool of ACs and HCs at this timepoint (Figure 6D). This was further confirmed by the elevated expression of both *ONECUT2* and *ONECUT1*, which both regulate early born cell fates and are critical for HC genesis,^57,58^ and *GAD2, GLYTL1A, MEIS2*, and *STMN2*, which are linked to AC and HC fates^59–62^ (Figure 6D). One of surprising finding was the early upregulation of *LHX3* post-PF treatment at D45 and, to a lesser extent, D60 (Figure 6D). *LHX3* in rodent species has been linked to BC differentiation;^63–65^ however, we hypothesize that *LHX3* may be a transiently expressed marker of early retinal cell fates, such as cone PRs. Finally, we analyzed cellular maturation through using synaptic markers.^66^ We observed significant upregulation of a number of PR and pan-retina associated synaptic genes (e.g. *SV2*, *SNAP25, SYP, RIMS2*,etc.), especially following D45 and D60 treatments, with smaller changes in D90 treated organoids (Figure 6E).

To confirm if these changes were significant, we carried out differential expression analysis using DESeq2 analysis in R. Differential expression analysis confirmed our IHC data (Figure 6F). At all three time points (D45, D60, and D90), we observed a significant reduction in Notch targets and RPC genes. D45 treated cells had the largest and most significant increase in PR associated genes. D90 had fewer significantly different genes and mostly enriched in PR maturation genes. Gene Ontology (GO) analysis on the RNAseq data showed that the most significant effect of PF-treatment on retinal organoids was a downregulation of cell cycle, mitosis, and DNA replication, which in turn resulted in a significant increase in gene networks associated with visual perception and synaptic organization and activity (Supplementary Figure S4).

## DISCUSSION

In this study, we describe a methodology to exploit human retinal progenitor competency states in retinal organoids to drive enriched PR populations. We show that the Notch pathway is active in RPCs within hPSC-derived retinal organoids, and that this pathway can be effectively inhibited using the Notch inhibitor, PF-30184014. Using this approach, we generated highly enriched cultures of PRs with significantly reduced numbers of proliferating retinal stem cells. The broader goal of this work is to generate a more uniform source of PRs that can be used for cell replacement therapies to aid in visual recovery.

The most unanticipated and interesting finding in our studies was the PR bias in our synchronized differentiated retinal organoids. When treated early (D45 or D60), we observed a significant rise in the cone PR population, shown via both IHC and bulk RNAseq. The same phenomenon occurred with the rod PR cell population when organoids were treated at D90, though to a smaller extent. While we expected a noticeable increase in other inner retinal neurons after PF-treatment, our data showed no significant differences to inner retinal neuron markers in PF treated organoids, even at the D60 treatment timepoint group. These results do, however, align well with previous studies showing that *Notch1* plays a role in the inhibition of cone and rod PR fate.^11,67,68^ Conditional genetic knockdown of Notch in early RPCs in mice has been shown to result in cone over-production at the expense of other retinal neurons, while a later-onset knockdown results in rod over-production. Recent studies using well-defined cis-regulatory elements for various progenitor states, including VSX2 for multipotent RPCs and THRB for cone-restricted RPCs, to analyze the effects of Notch signaling inhibition suggest that Notch regulates the formation of restricted RPC states from multipotent RPCs.^69^ This could be the potential mechanism driving specific fate decisions in human retina development. Additionally, Notch activity has been shown to be critical for the final MG fate specification.^52–54^ Persistent Notch activity promotes expression of downstream MG genes and stabilizes MG fate.^70^ Consistent with these studies, PF-treated retinal organoids show a paucity of MG cells.

Another interesting aspect of our studies was a lack of complete maturation of the differentiated cone and rod PRs. While these cells expressed early cone and rod differentiation markers, including RCVRN and ARR3, the more mature markers, such as the opsins, were lacking in these prematurely differentiated cells despite long-term culture for 70- and 140-days post-treatment. Similar observations have been observed in Zebrafish Notch knockdown studies.^54^ Together, these data suggest that other native retina cell types may be critical for driving full maturation of PRs, either directly or through secreted factors that may be missing in our PF-treated organoids. Recently, a number of factors have been shown to induce PR maturation and opsin expression in retinal organoids, including 9-cis, DHA, FGF-1, and thyroid hormone T3.^71–73^ It remains to be seen if addition of these factors could potentially drive maturation in PF-treated organoids. Along with a lack of maturation, there was an obvious loss of lamination and impaired organization in the PF-treated organoids. We hypothesize that this is due to the absence of critical cues from RPCs and/or MG, which likely provide spatial information to newly differentiated neurons. This disorganization also likely contributes to the lack of PR maturation, including the development of inner and outer segments. That being said, one advantage of generating an enriched pool of immature PRs is that these cells will likely achieve better integration following transplantation, compared to more mature PRs.^74,75^

In summary, the results reported here show that Notch signaling is necessary for RPC maintenance in hPSC-derived retinal organoids. Small-molecule Notch pathway inhibition using a compound validated in human studies, PF-03084014, can drive PR specific synchronized differentiation in retinal organoids to generate enriched cultures of immature cone and rod PRs which could be useful for cell replacement approaches to treat in severe PR degenerative disorders in affected patients.

## EXPERIMENTAL PROCEDURES

### Cell culture

The hESC line (CRXp-GFP H9) was kindly provided by Dr. Anand Swaroop. The iPSC line was kindly provided by Dr. Xianmin Zeng. Cells were maintained on Matrigel (Corning 354234) coated plates in mTeSR1 medium (StemCell Technologies) or mTeSR Plus medium (StemCell Technologies) in a 5% CO2/5% O2 incubator and passaged every 3-4 days using EDTA.

### Retinal organoid differentiation

Retinal organoids were differentiated via the embryoid body (EB) approach as described previously (Chirco et al., 2021).

### Notch inhibition

PF-30184014 (PF; Sigma Aldrich PZ0298-5MG) was prepared by dissolving it in DEPC water to a working concentration of 1mM. Aliquots were stored at −80°C until use. On treatment day, organoids were given their regular media with 10μM PF added. After 3 days, fresh non-PF media was given every 2-3 days until collection day. The reconstituted PF loses activity in about 3 months even at −80°C, so care was taken to use fresh compound.

### Bulk RNAseq data analysis

Control and PF-treated organoids were collected 28-days post-treatment for mRNA extraction as above for RNAseq analysis. Library preparation and Illumina-based transcriptome sequencing was carried out at Novogene. Following QC (error rates <0.03%), alignments were parsed using STAR (v2.5) program^76^ and mapped to Homo sapiens genome assembly GRCh37 (hg19). Heatmaps of key genes was graph using BEAVR web package (v1.0.10)^77^ in Docker. Differential expressions were determined through DESeq2 R package (v1.14.1).^78^ Genes with an adjusted p-value <0.05 found by DESeq2 were assigned as differentially expressed. Data was graphed in R using Enhanced Volcano plots package.^79^ Gene Ontology (GO) enrichment analysis of differentially expressed genes was carried out by ClusterProfiler R package (v2.4.3),^80^ in which gene length bias was corrected.

## Data availability

RNAseq data will be freely available at NCBI GEO at the time of publication.

## Acknowledgements

We would like to thank the members of the Lamba lab for helpful discussions and suggestions. The research presented here is supported by the National Eye Institute (U24 EY029891 and R01 EY032197 to DAL; P30 (EY002162) Vision Core grant to UCSF Dept of Ophthalmology), and the Research to Prevent Blindness (unrestricted grant to UCSF Dept of Ophthalmology).

We would like to thank Suling Wang for her assistance in creating Figure 1A.

## Supplementary Information

**Table S1.** Primary Antibodies for immunofluorescence staining.

| **Target** | **Host** | **Manufacturer** | **Catalog #** | **Dilution** |
| --- | --- | --- | --- | --- |
| **ARR3** | Mouse | Peter MacLeish Lab | n/a | 1:200 |
| **ARR3 (7G6)** | Mouse | Sigma Aldrich | MABN2636 | 1:500 |
| **BRN3A (14A6)** | Mouse | SCBT | sc-8429 | 1:100 |
| **CRX** | Rabbit | Abcam | ab140603 | 1:250 |
| **Glutamine Synthetase** | Rabbit | Abcam | ab49873 | 1:5000 |
| **HuC/D** | Mouse | Invitrogen | A21271 | 1:250 |
| **ISLET-1/2** | Mouse | DSHB | 39.4D5 | 1:5 |
| **KI67** | Rabbit | Abcam | ab15580 | 1:250 |
| **LHX1** | Mouse | DSHB | 4F2-s | 1:50 |
| **LHX2** | Mouse | DSHB | PCRP-LHX2-2G5 | 1:100 |
| **M/L-opsin** | Rabbit | Millipore Sigma | AB5405 | 1:2000 |
| **NRL** | Goat | R&D Systems | AF2945 | 1:200 |
| **OTX2** | Goat | R&D Systems | BAF1979 | 1:400 |
| **PAX6** | Mouse | DSHB | PAX6-s | 1:100 |
| **Recoverin** | Rabbit | EMD Millipore | AB5585 | 1:1000 |
| **Rhodopsin (4D2)** | Mouse | Millipore Sigma | MABN15 | 1:2000 |
| **RXRγ (Y-20)** | Rabbit | SCBT | sc-555 | 1:200 |
| **S-opsin** | Rabbit | Millipore Sigma | AB5407 | 1:2000 |
| **SAG (E-3)** | Mouse | SCBT | sc-166383 | 1:250 |
| **VSX2 (E-12)** | Mouse | SCBT | sc-365519 | 1:100 |

**Table S2.** Secondary antibodies for immunofluorescence staining.

| **Target** | **Host** | **Manufacturer** | **Catalog #** | **Dilution** |
| --- | --- | --- | --- | --- |
| **Anti-Mouse (488 Alexa Fluor)** | Donkey | Invitrogen | A-21202 | 1:500 |
| **Anti-Mouse (555 Alexa Fluor)** | Donkey | Invitrogen | A-31570 | 1:500 |
| **Anti-Mouse (647 Alexa Fluor)** | Donkey | Invitrogen | A-31571 | 1:500 |
| **Anti-Goat (488 Alexa Fluor)** | Donkey | Invitrogen | A-11055 | 1:500 |
| **Anti-Goat (555 Alexa Fluor)** | Donkey | Invitrogen | A-21432 | 1:500 |
| **Anti-Goat (647 Alexa Fluor)** | Donkey | Invitrogen | A-21447 | 1:500 |
| **Anti-Rabbit (488 Alexa Fluor)** | Donkey | Invitrogen | A-21206 | 1:500 |
| **Anti-Rabbit (555 Alexa Fluor)** | Donkey | Invitrogen | A-31572 | 1:500 |
| **Anti-Rabbit (647 Alexa Fluor)** | Donkey | Invitrogen | A-31573 | 1:500 |

**Table S3.**
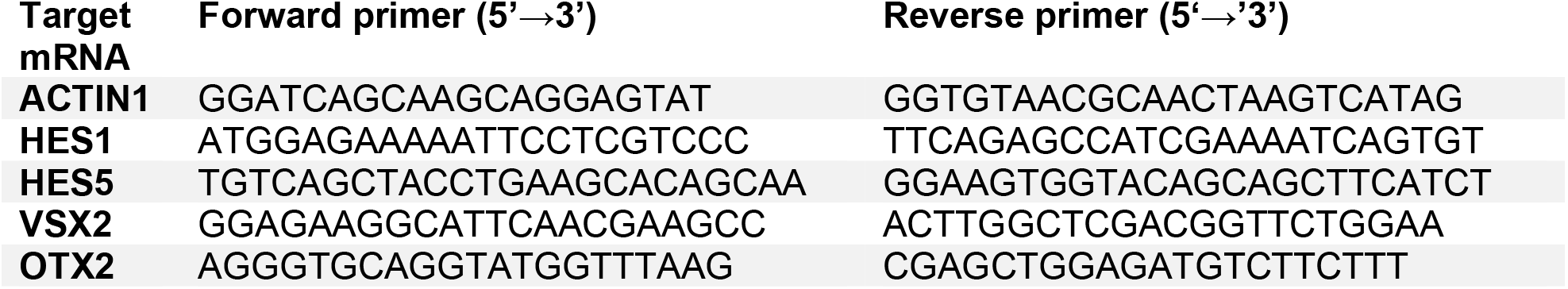
qPCR and RT-PCR Primers.

### Supplemental Experimental Procedures

#### hPSC passaging

For passaging, cultures were washed twice with DPBS (Corning) and incubated at room temperature for 3 minutes with 500μM EDTA in DPBS. The solution was carefully removed, and cells were gently rinsed with fresh DPBS. Fresh mTeSR1 or mTeSR Plus medium was added to the well and cells were lifted using a cell scraper. Cell colonies were broken into small clumps and passaged approximately 1:10 into a fresh Matrigel-coated well.

#### Retinal organoid differentiation

Briefly, hPSC colonies were lifted using Dispase and transferred to a 6-well suspension plate. EBs were gradually transitioned from mTeSR1 or mTeSR Plus to Neural Induction Medium (NIM) from D0 to D3. On D6, fresh NIM was supplemented with 1.5nM BMP4 (PeproTech 120-05ET). On D7, the EBs were transferred onto a Matrigel-coated plate to allow EBs to adhere.

The concentration of BMP4 was gradually reduced via half media changes every other day from D7 to D15. From D16-D29, the organoids were cultured in Retinal Differentiation Medium (RDM). Between D25-28, regions with clear retina-like morphology were gently lifted using a cell scraper, broken up by pipetting with a 5ml serological pipette 2-3 times, and transferred to a suspension culture plate. After an additional 24-48 hours, early retinal organoids can be identified by the translucent outer edge and transferred to a fresh suspension well as needed. From D30 onwards, the developing retinal organoids were cultured in 3D-RDM medium supplemented with 1μM all-trans retinoic acid (Millipore Sigma R2625), which was discontinued after D120.

##### Media formulations for retinal differentiation protocol

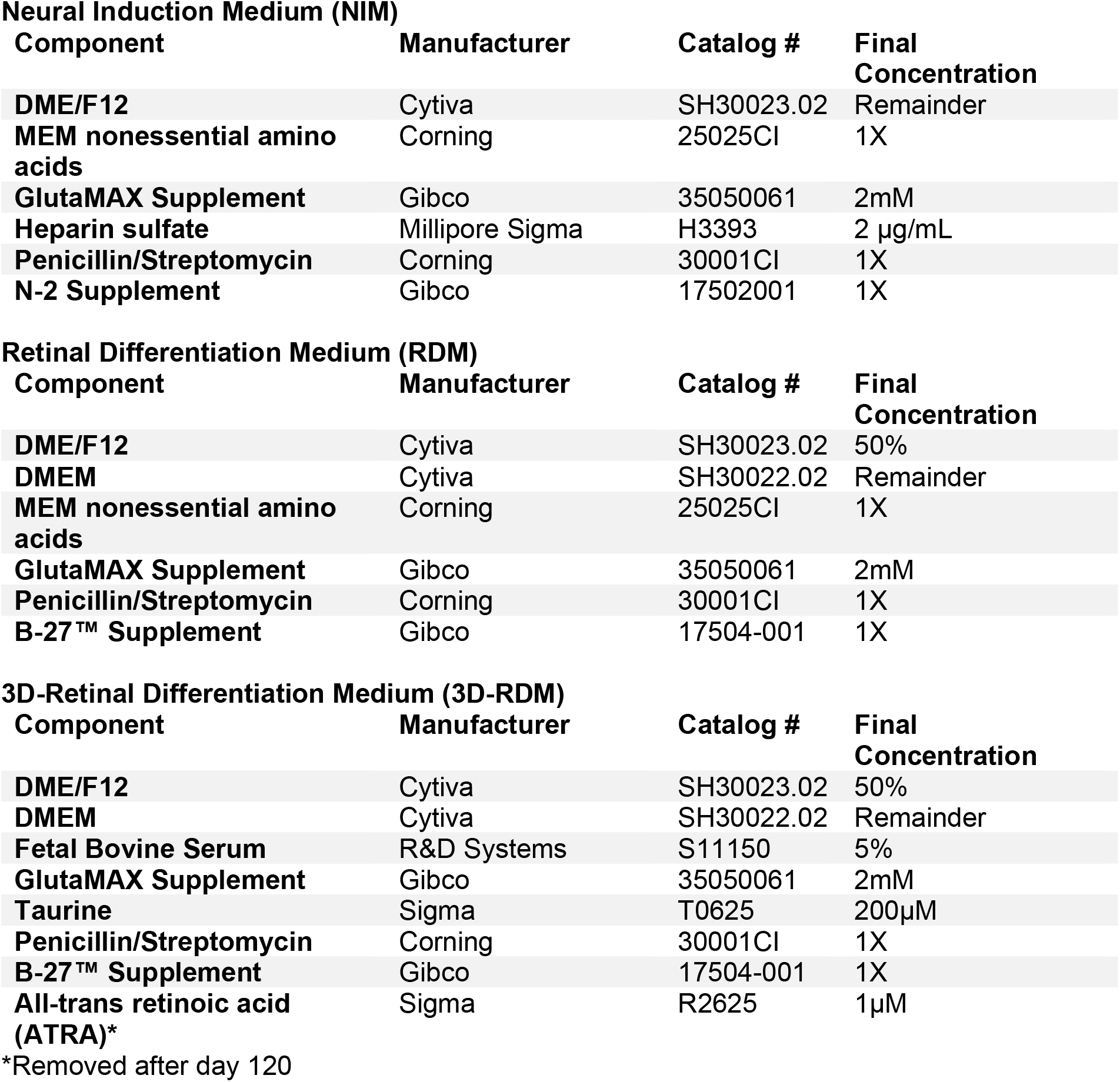

**Neural Induction Medium (NIM)**
| Component | Manufacturer | Catalog # | Final Concentration |
| --- | --- | --- | --- |
| DME/F12 | Cytiva | SH30023.02 | Remainder |
| MEM nonessential amino acids | Corning | 25025CI | 1X |
| GlutaMAX Supplement | Gibco | 35050061 | 2mM |
| Heparin sulfate | Millipore Sigma | H3393 | 2 µg/mL |
| Penicillin/Streptomycin | Corning | 30001CI | 1X |
| N-2 Supplement | Gibco | 17502001 | 1X |

**Retinal Differentiation Medium (RDM)**
| Component | Manufacturer | Catalog # | Final Concentration |
| --- | --- | --- | --- |
| DME/F12 | Cytiva | SH30023.02 | 50% |
| DMEM | Cytiva | SH30022.02 | Remainder |
| MEM nonessential amino acids | Corning | 25025CI | 1X |
| GlutaMAX Supplement | Gibco | 35050061 | 2mM |
| Penicillin/Streptomycin | Corning | 30001CI | 1X |
| B-27™ Supplement | Gibco | 17504-001 | 1X |

**3D-Retinal Differentiation Medium (3D-RDM)**
| Component | Manufacturer | Catalog # | Final Concentration |
| --- | --- | --- | --- |
| DME/F12 | Cytiva | SH30023.02 | 50% |
| DMEM | Cytiva | SH30022.02 | Remainder |
| Fetal Bovine Serum | R&D Systems | S11150 | 5% |
| GlutaMAX Supplement | Gibco | 35050061 | 2mM |
| Taurine | Sigma | T0625 | 200µM |
| Penicillin/Streptomycin | Corning | 30001CI | 1X |
| B-27™ Supplement | Gibco | 17504-001 | 1X |
| All-trans retinoic acid (ATRA)* | Sigma | R2625 | 1µM |

\*Removed after day 120

#### Histology and tissue immunostaining

Organoids were transferred from the cell culture dish into a 1.7ml tube and quickly washed with PBS. Approximately 500μl 4% PFA in PBS was added and the tube was left to incubate on ice for 25 minutes. The organoids were washed twice with PBS for 5 minutes at room temperature and then put through a sucrose series (15% and 30% in PBS) until the organoids sank to the bottom of the tube. The organoids were embedded in a 2:1 solution of 20% sucrose/OCT. Cryoblocks were stored in the −80°C at least overnight prior to sectioning. 7μm-thick sections were collected on SuperFrost slides (VWR 48311-703) and stored at −80°C until staining.

Prior to immunostaining, slides were warmed to room temperature. Sections were outlined with Pap pen (EMS 71310) and allowed to dry for 20 minutes. Slides were then washed with PBS for 5 minutes followed by blocking with 10% NDS (Millipore S30-100ML) for 1 hour at room temperature in humidified chamber. Sections were then incubated with primary antibodies overnight at 4°C (refer to supplementary for antibody list). The next day, slides were washed in PBS 3 times for 5-10 minutes each at room temperature and then incubated with secondary antibodies for 1 hour at room temperature (refer to supplementary for antibody list). Secondary solution was tapped off the slides and DAPI (0.5μg/ml, Roche 10236276001) was applied for 5 minutes. Slides were washed with PBS three times for 5-10 minutes each and coverslips were applied using Fluoromount-G (EMS 17984-25). Images were taken using a LSM700 confocal with a 40X oil objective.

#### RNA extraction

About 3-5 organoids were transferred from the cell culture dish to a 1.7ml tube and quickly washed once with sterile PBS. After aspirating the PBS, 50μl TriReagent (Zymo R2050-1-200) was added. Tubes were frozen at −80°C until time of extraction. Samples were thawed on ice and 250μl of fresh TriReagent was added. Organoids were homogenized with a handheld electronic pestle mixer. At room temperature, 60μl chloroform was added to the tube and the solution was gently vortexed for 10 seconds. The solution was incubated at room temperature for 2 minutes and then centrifuged at 16.1xG at 4°C for 15 minutes. The top aqueous layer was transferred to a separate tube and GlycoBlue (1:100, Invitrogen AM9515) was added. Isopropanol (1 volume) was added and then the tube was gently vortexed for 10 seconds. Solution was then incubated at −20°C overnight before being spun at 16.1xG at 4°C for 15 minutes. The supernatant was then discarded once a blue pellet was identified. The pellet was washed with 1mL of cold 70% EtOH and centrifuged again at 16.1xG at 4°C for 15 minutes. This step was repeated. The solution was decanted, and the remaining pellet was allowed to air dry at room temperature for about 15 minutes before being dissolved in DEPC water and incubated at 55°C for 5 minutes. RNA was spec’d using a nanodrop. RNA (≤4μg) was cleaned using the TURBO DNA-free™ Kit (Invitrogen AM1907) and following manufacturer’s instructions.

#### qPCR analysis

cDNA was generated using the iScript cDNA Synthesis Kit (Bio-Rad 1708891). To run qPCR experiments, the iTaq Universal SYBR Green Supermix (Bio-Rad 1725124) mixed with specific forward and reverse primer sequences (see Supplementary Table S3), cDNA, and water to a final reaction volume of 10μl. The thermocycling parameters were as follows: 95°C for 30 seconds, followed by amplification for 40 cycles at 95°C for 5 seconds and at 60°C for 25 seconds and was followed by melting curve analysis from 60°C to 95°C. All data summary graphs were made using GraphPad Prism 9, and all statistical analyses were performed via two-way ANOVA analysis with multiple comparisons.

## References

1. Lamba DA, Karl MO, Ware CB, Reh TA. Efficient generation of retinal progenitor cells from human embryonic stem cells. Proc Natl Acad Sci U S A. 2006;103(34):12769–12774. doi:10.1073/pnas.0601990103

2. Zhu J, Reynolds J, Garcia T, et al. Generation of Transplantable Retinal Photoreceptors from a Current Good Manufacturing Practice-Manufactured Human Induced Pluripotent Stem Cell Line: Transplantable Photoreceptors from cGMP iPSCs. STEM CELLS Transl Med. 2018;7(2):210–219. doi:10.1002/sctm.17-0205

3. Chirco KR, Chew S, Moore AT, Duncan JL, Lamba DA. Allele-specific gene editing to rescue dominant CRX-associated LCA7 phenotypes in a retinal organoid model. Stem Cell Rep. 2021;16(11):2690–2702. doi:10.1016/j.stemcr.2021.09.007

4. Ohlemacher SK, Sridhar A, Xiao Y, et al. Stepwise Differentiation of Retinal Ganglion Cells from Human Pluripotent Stem Cells Enables Analysis of Glaucomatous Neurodegeneration: hPSC-derived RGCs and Glaucoma. STEM CELLS. 2016;34(6):1553–1562. doi:10.1002/stem.2356

5. Zhu J, Cifuentes H, Reynolds J, Lamba DA. Immunosuppression via Loss of IL2rγ Enhances Long-Term Functional Integration of hESC-Derived Photoreceptors in the Mouse Retina. Cell Stem Cell. 2017;20(3):374–384.e5. doi:10.1016/j.stem.2016.11.019

6. Gagliardi G, Ben M’Barek K, Chaffiol A, et al. Characterization and Transplantation of CD73-Positive Photoreceptors Isolated from Human iPSC-Derived Retinal Organoids. Stem Cell Rep. 2018;11(3):665–680. doi:10.1016/j.stemcr.2018.07.005

7. Lin B, McLelland B, Xue Y, et al. hESC-derived retina organoids produced by a scalable cGMP compatible process improve visual function after transplantation to immunodeficient RD rats. Invest Ophthalmol Vis Sci. 2020;61(7):2505.

8. Shirai H, Mandai M, Matsushita K, et al. Transplantation of human embryonic stem cell-derived retinal tissue in two primate models of retinal degeneration. Proc Natl Acad Sci U S A. 2016;113(1):E81–90. doi:10.1073/pnas.1512590113

9. Arai MA, Akamine R, Tsuchiya A, et al. The Notch inhibitor cowanin accelerates nicastrin degradation. Sci Rep. 2018;8(1):5376. doi:10.1038/s41598-018-23698-4

10. Bray SJ. Notch signalling: a simple pathway becomes complex. Nat Rev Mol Cell Biol. 2006;7(9):678–689. doi:10.1038/nrm2009

11. Jadhav AP, Cho SH, Cepko CL. Notch activity permits retinal cells to progress through multiple progenitor states and acquire a stem cell property. Proc Natl Acad Sci. 2006; 103(50):18998–19003. doi:10.1073/pnas.0608155103

12. Nelson BR, Hartman BH, Georgi SA, Lan MS, Reh TA. Transient inactivation of Notch signaling synchronizes differentiation of neural progenitor cells. Dev Biol. 2007;304(2):479–498. doi:10.1016/j.ydbio.2007.01.001

13. Hayes S, Nelson BR, Buckingham B, Reh TA. Notch signaling regulates regeneration in the avian retina. Dev Biol. 2007;312(1):300–311. doi:10.1016/j.ydbio.2007.09.046

14. Raymond PA, Barthel LK, Bernardos RL, Perkowski JJ. Molecular characterization of retinal stem cells and their niches in adult zebrafish. BMC Dev Biol. 2006;6:36. doi:10.1186/1471-213X-6-36

15. Curigliano G, Aftimos PG, Dees EC, et al. Phase I dose-finding study of the gamma secretase inhibitor PF-03084014 (PF-4014) in combination with docetaxel in patients (pts) with advanced triple-negative breast cancer (TNBC). J Clin Oncol. 2015;33(15_suppl):1068–1068. doi:10.1200/jco.2015.33.15_suppl.1068

16. Kaufman ML, Park KU, Goodson NB, et al. Transcriptional profiling of murine retinas undergoing semi-synchronous cone photoreceptor differentiation. Dev Biol. 2019;453(2):155–167. doi:10.1016/j.ydbio.2019.05.016

17. Pei Y, Sierra G, Sivapatham R, Swistowski A, Rao MS, Zeng X. A platform for rapid generation of single and multiplexed reporters in human iPSC lines. Sci Rep. 2015;5(1):9205. doi:10.1038/srep09205

18. Kaewkhaw R, Kaya KD, Brooks M, et al. Transcriptome Dynamics of Developing Photoreceptors in Three-Dimensional Retina Cultures Recapitulates Temporal Sequence of Human Cone and Rod Differentiation Revealing Cell Surface Markers and Gene Networks: Transcriptome of Developing Human Photoreceptors. STEM CELLS. 2015;33(12): 3504–3518. doi:10.1002/stem.2122

19. Ohtsuka T, Ishibashi M, Gradwohl G, Nakanishi S, Guillemot F, Kageyama R. Hes1 and Hes5 as notch effectors in mammalian neuronal differentiation. EMBO J. 1999;18(8):2196–2207. doi:10.1093/emboj/18.8.2196

20. Livne-Bar I, Pacal M, Cheung MC, et al. Chx10 is required to block photoreceptor differentiation but is dispensable for progenitor proliferation in the postnatal retina. Proc Natl Acad Sci U S A. 2006; 103(13):4988–4993. doi:10.1073/pnas.0600083103

21. Vitorino M, Jusuf PR, Maurus D, Kimura Y, Higashijima S ichi, Harris WA. Vsx2 in the zebrafish retina: restricted lineages through derepression. Neural Develop. 2009;4(1):14. doi:10.1186/1749-8104-4-14

22. Glubrecht DD, Kim JH, Russell L, Bamforth JS, Godbout R. Differential CRX and OTX2 expression in human retina and retinoblastoma. J Neurochem. 2009;111(1):250–263. doi:10.1111/j.1471-4159.2009.06322.x

23. Yamamoto H, Kon T, Omori Y, Furukawa T. Functional and Evolutionary Diversification of Otx2 and Crx in Vertebrate Retinal Photoreceptor and Bipolar Cell Development. Cell Rep. 2020;30(3):658–671.e5. doi:10.1016/j.celrep.2019.12.072

24. Ruzycki PA, Zhang X, Chen S. CRX directs photoreceptor differentiation by accelerating chromatin remodeling at specific target sites. Epigenetics Chromatin. 2018;11(1):42. doi:10.1186/s13072-018-0212-2

25. Pacal M, Bremner R. Mapping differentiation kinetics in the mouse retina reveals an extensive period of cell cycle protein expression in post-mitotic newborn neurons. Dev Dyn. 2012;241(10):1525–1544. doi:10.1002/dvdy.23840

26. Marquardt T, Ashery-Padan R, Andrejewski N, Scardigli R, Guillemot F, Gruss P. Pax6 Is Required for the Multipotent State of Retinal Progenitor Cells. Cell. 2001;105(1):43–55. doi:10.1016/S0092-8674(01)00295-1

27. Nishina S, Kohsaka S, Yamaguchi Y, et al. PAX6 expression in the developing human eye. Br J Ophthalmol. 1999;83(6):723–727.

28. Shaham O, Menuchin Y, Farhy C, Ashery-Padan R. Pax6: A multi-level regulator of ocular development. Prog Retin Eye Res. 2012;31(5):351–376. doi:10.1016/j.preteyeres.2012.04.002

29. de Melo J, Zibetti C, Clark BS, et al. Lhx2 Is an Essential Factor for Retinal Gliogenesis and Notch Signaling. J Neurosci Off J Soc Neurosci. 2016;36(8):2391–2405. doi:10.1523/JNEUROSCI.3145-15.2016

30. Bassett EA, Wallace VA. Cell fate determination in the vertebrate retina. Trends Neurosci. 2012;35(9):565–573. doi:10.1016/j.tins.2012.05.004

31. Brzezinski JA, Reh TA. Photoreceptor cell fate specification in vertebrates. Development. 2015;142(19):3263–3273. doi:10.1242/dev.127043

32. Cepko C. Intrinsically different retinal progenitor cells produce specific types of progeny. Nat Rev Neurosci. 2014;15(9):615–627. doi:10.1038/nrn3767

33. Xiang M. Intrinsic control of mammalian retinogenesis. Cell Mol Life Sci. 2013;70(14):2519–2532. doi:10.1007/s00018-012-1183-2

34. Hoshino A, Ratnapriya R, Brooks MJ, et al. Molecular Anatomy of the Developing Human Retina. Dev Cell. 2017;43(6):763–779.e4. doi:10.1016/j.devcel.2017.10.029

35. Sridhar A, Hoshino A, Finkbeiner CR, et al. Single-Cell Transcriptomic Comparison of Human Fetal Retina, hPSC-Derived Retinal Organoids, and Long-Term Retinal Cultures. Cell Rep. 2020;30(5):1644–1659.e4. doi:10.1016/j.celrep.2020.01.007

36. Boije H, MacDonald RB, Harris WA. Reconciling competence and transcriptional hierarchies with stochasticity in retinal lineages. Curr Opin Neurobiol. 2014;27:68–74. doi:10.1016/j.conb.2014.02.014

37. Bazhin AV, Schadendorf D, Philippov PP, Eichmüller SB. Recoverin as a cancer-retina antigen. Cancer Immunol Immunother CII. 2007;56(1):110–116. doi:10.1007/s00262-006-0132-z

38. Makino CL, Dodd RL, Chen J, et al. Recoverin Regulates Light-dependent Phosphodiesterase Activity in Retinal Rods. J Gen Physiol. 2004;123(6):729–741. doi:10.1085/jgp.200308994

39. Roberts MR, Hendrickson A, McGuire CR, Reh TA. Retinoid X receptor (gamma) is necessary to establish the S-opsin gradient in cone photoreceptors of the developing mouse retina. Invest Ophthalmol Vis Sci. 2005;46(8):2897–2904. doi:10.1167/iovs.05-0093

40. Sakuma H, Murakami A, Fujimaki T, Inana G. Isolation and characterization of the human X-arrestin gene. Gene. 1998;224(1):87–95. doi:10.1016/S0378-1119(98)00510-1

41. Mears AJ, Kondo M, Swain PK, et al. Nrl is required for rod photoreceptor development. Nat Genet. 2001;29(4):447–452. doi:10.1038/ng774

42. Boije H, Shirazi Fard S, Edqvist PH, Hallböök F. Horizontal Cells, the Odd Ones Out in the Retina, Give Insights into Development and Disease. Front Neuroanat. 2016;10. Accessed January 19, 2022. https://www.frontiersin.org/article/10.3389/fnana.2016.00077

43. Liu W, Wang JH, Xiang M. Specific expression of the LIM/Homeodomain protein Lim-1 in horizontal cells during retinogenesis. Dev Dyn. 2000;217(3):320–325. doi:10.1002/(SICI)1097-0177(200003)217:3<320::AID-DVDY10>3.0.CO;2-F

44. Badea TC, Cahill H, Ecker J, Hattar S, Nathans J. Distinct Roles of Transcription Factors Brn3a and Brn3b in Controlling the Development, Morphology, and Function of Retinal Ganglion Cells. Neuron. 2009;61(6):852–864. doi:10.1016/j.neuron.2009.01.020

45. Ekström P, Johansson K. Differentiation of ganglion cells and amacrine cells in the rat retina: correlation with expression of HuC/D and GAP-43 proteins. Dev Brain Res. 2003; 145(1):1–8. doi:10.1016/S0165-3806(03)00170-6

46. Elshatory Y, Everhart D, Deng M, Xie X, Barlow RB, Gan L. Islet-1 Controls the Differentiation of Retinal Bipolar and Cholinergic Amacrine Cells. J Neurosci. 2007;27(46):12707–12720. doi:10.1523/JNEUROSCI.3951-07.2007

47. Martín-Partido G, Francisco-Morcillo J. The role of Islet-1 in cell specification, differentiation, and maintenance of phenotypes in the vertebrate neural retina. Neural Regen Res. 2015;10(12):1951–1952. doi:10.4103/1673-5374.165301

48. Fischer AJ, Foster S, Scott MA, Sherwood P. The transient expression of LIM-domain transcription factors is coincident with the delayed maturation of photoreceptors in the chicken retina. J Comp Neurol. 2008;506(4):584–603. doi:10.1002/cne.21578

49. O’Brien KMB, Cheng H, Jiang Y, Schulte D, Swaroop A, Hendrickson AE. Expression of Photoreceptor-Specific Nuclear Receptor NR2E3 in Rod Photoreceptors of Fetal Human Retina. Invest Ophthalmol Vis Sci. 2004;45(8):2807–2812. doi:10.1167/iovs.03-1317

50. Akimoto M, Cheng H, Zhu D, et al. Targeting of GFP to newborn rods by Nrl promoter and temporal expression profiling of flow-sorted photoreceptors. Proc Natl Acad Sci. 2006; 103(10):3890–3895. doi:10.1073/pnas.0508214103

51. Peng GH, Ahmad O, Ahmad F, Liu J, Chen S. The photoreceptor-specific nuclear receptor Nr2e3 interacts with Crx and exerts opposing effects on the transcription of rod versus cone genes. Hum Mol Genet. 2005;14(6):747–764. doi:10.1093/hmg/ddi070

52. Furukawa T, Mukherjee S, Bao ZZ, Morrow EM, Cepko CL. rax, Hes1, and notch1 promote the formation of Müller glia by postnatal retinal progenitor cells. Neuron. 2000;26(2):383–394. doi:10.1016/s0896-6273(00)81171-x

53. Hojo M, Ohtsuka T, Hashimoto N, Gradwohl G, Guillemot F, Kageyama R. Glial cell fate specification modulated by the bHLH gene Hes5 in mouse retina. Dev Camb Engl. 2000;127(12):2515–2522. doi:10.1242/dev.127.12.2515

54. Bernardos RL, Lentz SI, Wolfe MS, Raymond PA. Notch-Delta signaling is required for spatial patterning and Müller glia differentiation in the zebrafish retina. Dev Biol. 2005;278(2):381–395. doi:10.1016/j.ydbio.2004.11.018

55. Linser P, Moscona AA. Induction of glutamine synthetase in embryonic neural retina: localization in Müller fibers and dependence on cell interactions. Proc Natl Acad Sci U S A. 1979;76(12):6476–6480. doi:10.1073/pnas.76.12.6476

56. Dyer MA, Livesey FJ, Cepko CL, Oliver G. Prox1 function controls progenitor cell proliferation and horizontal cell genesis in the mammalian retina. Nat Genet. 2003;34(1):53–58. doi:10.1038/ng1144

57. Klimova L, Antosova B, Kuzelova A, Strnad H, Kozmik Z. Onecut1 and Onecut2 transcription factors operate downstream of Pax6 to regulate horizontal cell development. Dev Biol. 2015;402(1):48–60. doi:10.1016/j.ydbio.2015.02.023

58. Sapkota D, Chintala H, Wu F, Fliesler SJ, Hu Z, Mu X. Onecut1 and Onecut2 redundantly regulate early retinal cell fates during development. Proc Natl Acad Sci U S A. 2014;111(39):E4086–4095. doi:10.1073/pnas.1405354111

59. Bumsted-O’Brien KM, Hendrickson A, Haverkamp S, Ashery-Padan R, Schulte D. Expression of the homeodomain transcription factor Meis2 in the embryonic and postnatal retina. J Comp Neurol. 2007;505(1):58–72. doi:10.1002/cne.21458

60. Crooks J, Kolb H. Localization of GABA, glycine, glutamate and tyrosine hydroxylase in the human retina. J Comp Neurol. 1992;315(3):287–302. doi:10.1002/cne.903150305

61. Lu Y, Shiau F, Yi W, et al. Single-Cell Analysis of Human Retina Identifies Evolutionarily Conserved and Species-Specific Mechanisms Controlling Development. Dev Cell. 2020;53(4):473–491.e9. doi:10.1016/j.devcel.2020.04.009

62. Nakazawa T, Nakano I, Furuyama T, Morii H, Tamai M, Mori N. The SCG10-related gene family in the developing rat retina: persistent expression of SCLIP and stathmin in mature ganglion cell layer. Brain Res. 2000;861(2):399–407. doi:10.1016/s0006-8993(00)02056-4

63. Kim Y, Lim S, Ha T, et al. The LIM protein complex establishes a retinal circuitry of visual adaptation by regulating Pax6 α-enhancer activity. eLife. 2017;6:e21303. doi:10.7554/eLife.21303

64. Buenaventura DF, Corseri A, Emerson MM. Identification of Genes With Enriched Expression in Early Developing Mouse Cone Photoreceptors. Invest Ophthalmol Vis Sci. 2019;60(8):2787–2799. doi:10.1167/iovs.19-26951

65. Dong X, Yang H, Zhou X, et al. LIM-Homeodomain Transcription Factor LHX4 Is Required for the Differentiation of Retinal Rod Bipolar Cells and OFF-Cone Bipolar Subtypes. Cell Rep. 2020;32(11):108144. doi:10.1016/j.celrep.2020.108144

66. Burger CA, Jiang D, Mackin RD, Samuel MA. Development and maintenance of vision’s first synapse. Dev Biol. 2021;476:218–239. doi:10.1016/j.ydbio.2021.04.001

67. Yaron O, Farhy C, Marquardt T, Applebury M, Ashery-Padan R. Notch1 functions to suppress cone-photoreceptor fate specification in the developing mouse retina. Dev Camb Engl. 2006;133(7):1367–1378. doi:10.1242/dev.02311

68. Mizeracka K, DeMaso CR, Cepko CL. Notch1 is required in newly postmitotic cells to inhibit the rod photoreceptor fate. Dev Camb Engl. 2013;140(15):3188–3197. doi:10.1242/dev.090696

69. Chen X, Emerson MM. Notch signaling represses cone photoreceptor formation through the regulation of retinal progenitor cell states. Sci Rep. 2021;11(1):14525. doi:10.1038/s41598-021-93692-w

70. Nelson BR, Ueki Y, Reardon S, et al. Genome-wide analysis of Müller glial differentiation reveals a requirement for Notch signaling in postmitotic cells to maintain the glial fate. PloS One. 2011;6(8):e22817. doi:10.1371/journal.pone.0022817

71. Kelley RA, Chen HY, Swaroop A, Li T. Accelerated Development of Rod Photoreceptors in Retinal Organoids Derived from Human Pluripotent Stem Cells by Supplementation with 9-cis Retinal. STAR Protoc. 2020;1(1):100033. doi:10.1016/j.xpro.2020.100033

72. Brooks MJ, Chen HY, Kelley RA, et al. Improved Retinal Organoid Differentiation by Modulating Signaling Pathways Revealed by Comparative Transcriptome Analyses with Development In Vivo. Stem Cell Rep. Published online October 2019:S2213671119303388. doi:10.1016/j.stemcr.2019.09.009

73. Eldred KC, Hadyniak SE, Hussey KA, et al. Thyroid hormone signaling specifies cone subtypes in human retinal organoids. Science. 2018;362(6411):eaau6348. doi:10.1126/science.aau6348

74. Garita-Hernandez M, Lampič M, Chaffiol A, et al. Restoration of visual function by transplantation of optogenetically engineered photoreceptors. Nat Commun. 2019;10(1):4524. doi:10.1038/s41467-019-12330-2

75. Gonzalez-Cordero A, Kruczek K, Naeem A, et al. Recapitulation of Human Retinal Development from Human Pluripotent Stem Cells Generates Transplantable Populations of Cone Photoreceptors. Stem Cell Rep. 2017;9(3):820–837. doi:10.1016/j.stemcr.2017.07.022

76. Dobin A, Davis CA, Schlesinger F, et al. STAR: ultrafast universal RNA-seq aligner. Bioinforma Oxf Engl. 2013;29(1):15–21. doi:10.1093/bioinformatics/bts635

77. Perampalam P, Dick FA. BEAVR: a browser-based tool for the exploration and visualization of RNA-seq data. BMC Bioinformatics. 2020;21(1):221. doi:10.1186/s12859-020-03549-8

78. Anders S, Huber W. Differential expression analysis for sequence count data. Genome Biol. 2010;11(10):R106. doi:10.1186/gb-2010-11-10-r106

79. Blighe K, Rana S, Lewis M. EnhancedVolcano: Publication-Ready Volcano Plots with Enhanced Colouring and Labeling.; 2021. Accessed January 27, 2022. https://github.com/kevinblighe/EnhancedVolcano

80. Yu G, Wang LG, Han Y, He QY. clusterProfiler: an R package for comparing biological themes among gene clusters. Omics J Integr Biol. 2012;16(5):284–287. doi:10.1089/omi.2011.0118

