## Supplementary data for "Timed Notch Inhibition drives Photoreceptor fate specification in Human Retinal Organoids"

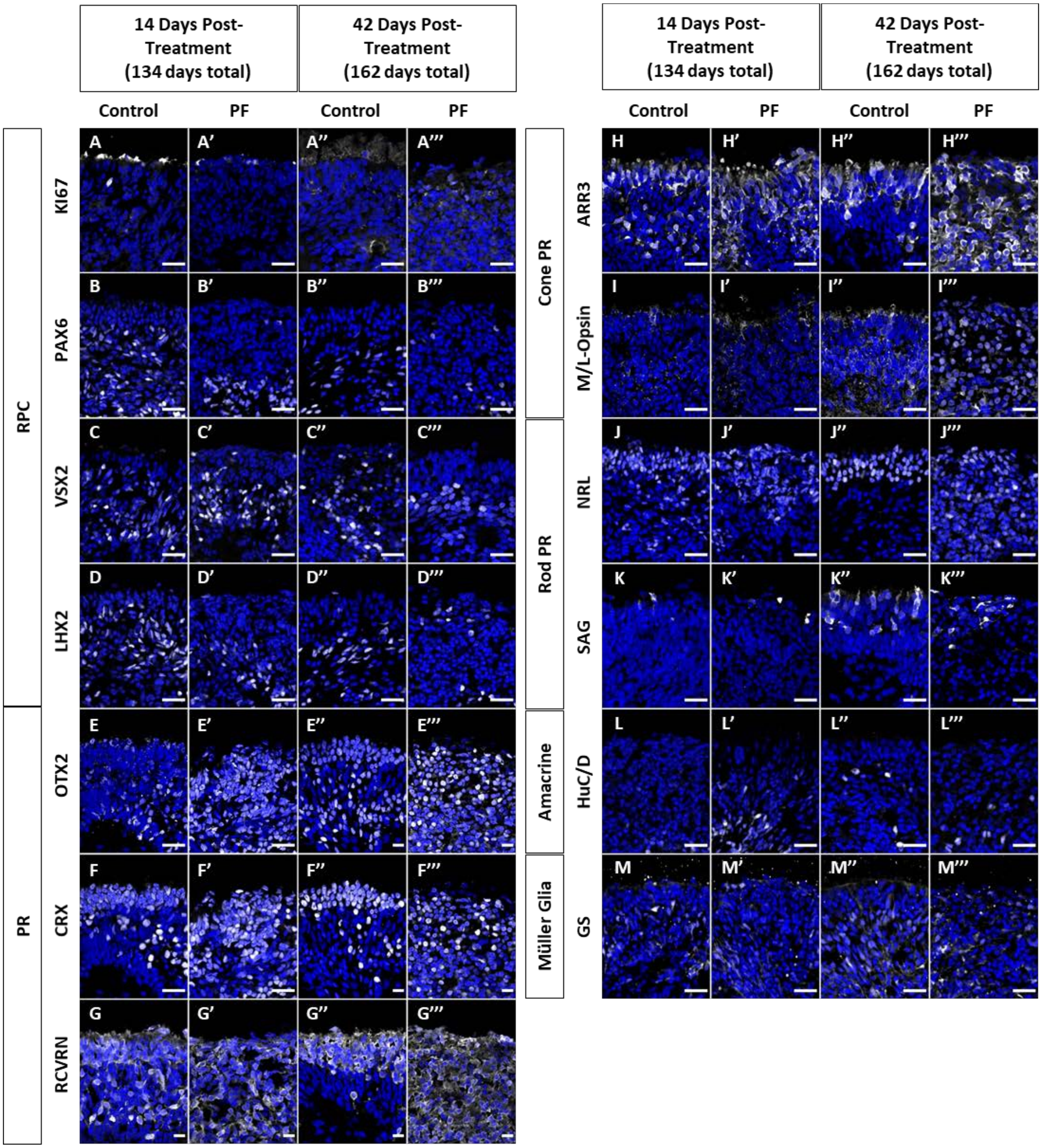


**Supplemental Figure 1. Notch inhibition at D120 causes little change retinal organoid differentiation after 14- and 42-days.** Immunofluorescence staining using antibodies against KI67 (A), PAX6 (B), VSX2 (C), LHX2 (D), OTX2 (E), CRX (F), RCVRN (G), ARR3 (H), M/L-Opsin (I), NRL (J), SAG (K), HuC/D (L), and GS (M) are shown in white. Markers are split into RPCs (A-D), pan photoreceptors (E-G), Cone PR (H-I), Rod PR (J-K), AC (L), and MG (M). Nuclei are counterstained with DAPI in blue. Scale bar, 25μm.

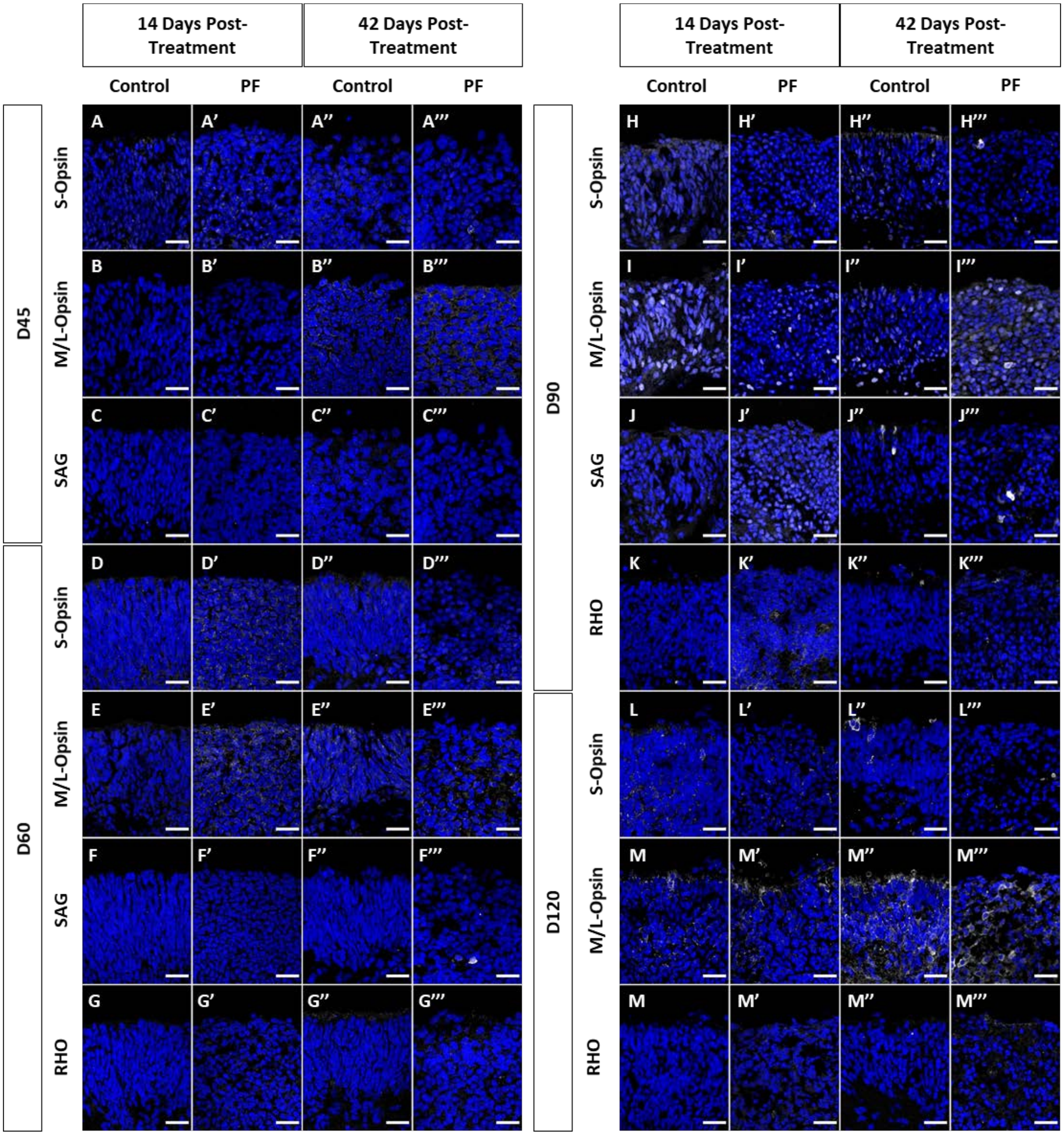


**Supplemental Figure 2. Notch inhibition causes reduced PR maturation in retinal organoids after 14- and 42-days.** Immunofluorescence staining using antibodies against S-Opsin (A, D, H, L), M/L-Opsin (B, E, I, M), SAG (C, F, J), and RHO (G, K, M) are shown in white for treatment groups D45 (A-C), D60 (D-G), D90 (H-K), and D120 (L-M). Nuclei are counterstained with DAPI in blue. Scale bar, 25μm.

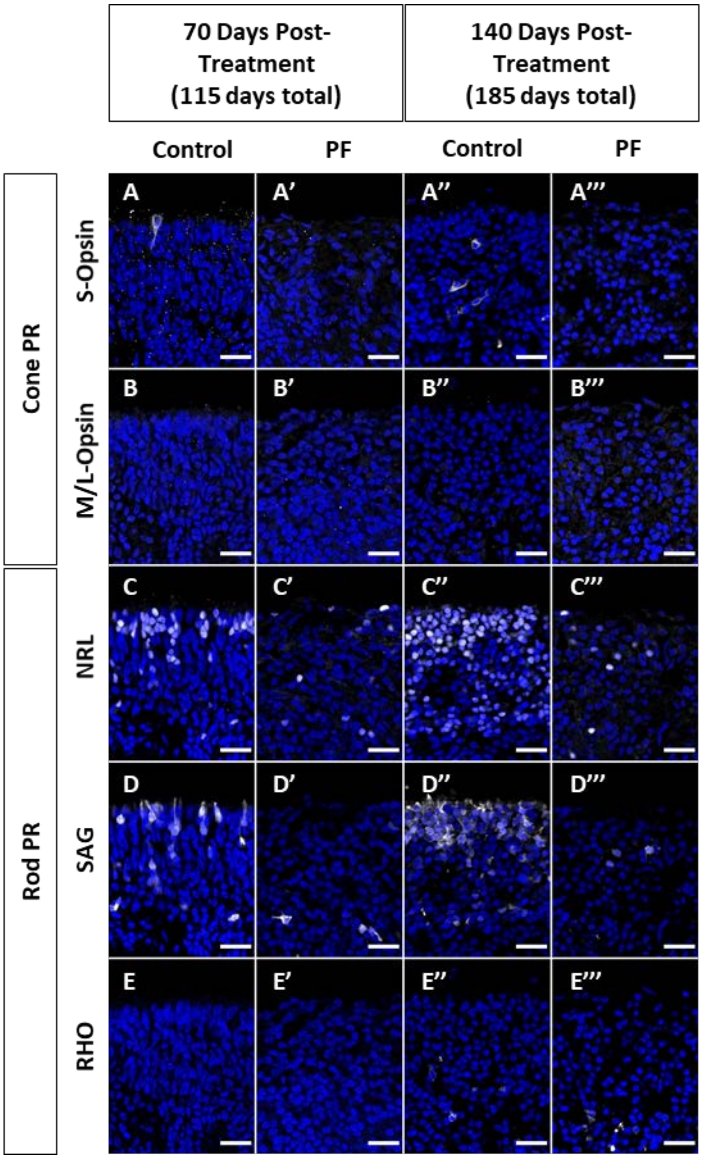


**Supplemental Figure 3. Notch inhibition causes reduced PR maturation in D45 PF-treated retinal organoids after 70- and 140-days.** Immunofluorescence staining using antibodies against S-Opsin (A), M/L-Opsin (B), NRL (C), SAG (D), and RHO (E) are shown in white. Markers are split into Cone PR (A-B) and Rod PR (C-E). Nuclei are counterstained with DAPI in blue. Scale bar, 25µm.

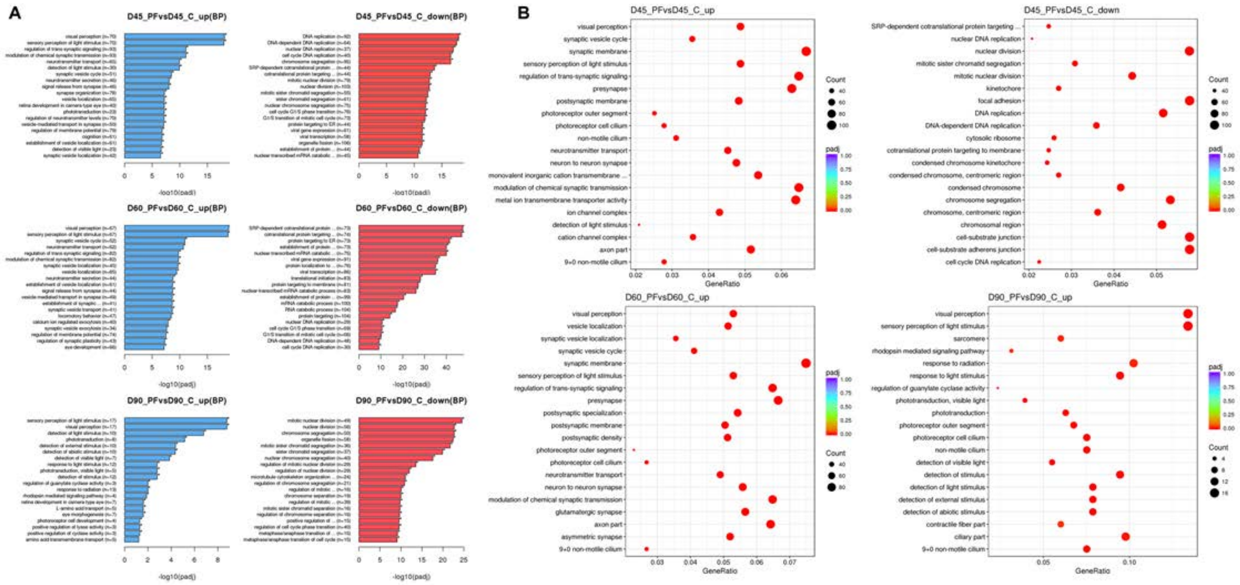


**Supplemental Figure 4. GO Analysis of differentially expressed genes following RNAseq analysis at all three time points.** (A) Bar graphs showing key Biological Processes either upregulated (blue) or downregulated (red) following PF treatment of D45, D60 and D90 organoids. (B) Dot Plots of enriched GO terms. The color of the dots represents the significance (p-value) for each enriched GO term. The size of the dots represents number of genes and Y-axis represents the enrichment signal strength as a percentage of genes included in the complete gene set.
